# SegColR: Deep Learning for Automated Segmentation and Color Extraction

**DOI:** 10.1101/2024.07.28.605475

**Authors:** James Boyko

## Abstract

Citizen science platforms like iNaturalist generate biodiversity data at an unprecedented scale, with observations on the order of hundreds of millions. However, extracting phenotypic information from these images, such as color of organisms, at such a large scale poses unique challenges for biologists. Some of the challenges are that manual extraction of phenotypic information can be subjective and time-consuming. Fortunately, with the maturation of computer vision and deep learning, there is an opportunity to automate large parts of the image processing pipeline. Here, I present SegColR, a user-friendly software package that leverages two state-of-the-art deep learning models - GroundingDINO and SegmentAnything - to enable automated segmentation and color extraction from images. The SegColR package provides an R-based interface, making it more accessible to evolutionary biologists and ecologists who may not have extensive coding experience. The SegColR pipeline allows users to load images, automatically segment them based on text prompts, and extract color information from the segmented regions. The package also includes visualization and data summarization functions to facilitate downstream analysis and interpretation of the results.

## 1 Introduction

Color is an important trait for many organisms, influencing various aspects of their ecology and physiology. It plays a crucial role across the tree of life in communication, pollination syndromes, mating behavior, predator-prey interactions, and thermal regulation (e.g., Endler 1993; Huyghe et al. 2007; Tsuchida et al. 2010; Finkbeiner et al. 2014; Mitchem 2017; Narbona et al. 2021; Cox and Davis Rabosky 2023). However, obtaining large-scale, high-quality color data can be a significant challenge for researchers. For example, while research museum collections house vast amounts of biodiversity information, the preservation techniques used may distort the original coloration of specimens (Pohland and Mullen 2006). Nonetheless, the ubiquity and importance of color in organismal biology means that accurately quantifying color traits is an important step towards understanding many aspects of ecology and evolution of organisms.

Citizen science initiatives like iNaturalist generate data in the form of crowdsourced observations and images, centralizing vast quantities of biodiversity data and providing opportunity to study color variation across the tree of life and at large spatial scales. Yet, the sheer scale of the data generated and the fact that it is typically not standardized, make it difficult to automate data collection and extraction. As such, most attempts to utilize resources like iNaturalist rely on manually extracting information image by image. Though this approach almost certainly results in high-quality datasets, for even relatively simple tasks like color quantification, extracting information from images will be time-consuming and likely require large teams of willing annotators. Furthermore, manual color extraction can be subject to measurement error, e.g. what one observer calls white another may call light green. Fortunately, the maturation of computer vision and deep learning techniques offers a promising solution to this challenge. By leveraging state-of-the-art machine learning models, it is now possible to automate large parts of the image processing pipeline, allowing biologists to extract phenotypic information from citizen science data at unprecedented scales.

Many of the most successful deep learning applications in recent years have been for computer vision tasks (LeCun et al. 2015; Goodfellow et al. 2016). Two key areas of computer vision that are particularly relevant for extracting color data from biodiversity images are object detection and instance segmentation. Object detection algorithms aim to locate and classify distinct objects within an image (Redmon et al. 2016; Ren et al. 2017), while instance segmentation models, delineate pixel-level boundaries of each instance of an object (He et al. 2017; Kirillov et al. 2020). By combining these capabilities, it becomes possible to focus color analysis only on the specific organisms of interest, rather than the entire scene. However, training a specialized computer vision model from scratch would require large, annotated datasets. This would be prohibitively time-consuming to collect, especially for diverse taxonomic groups. Fortunately, progress in computer vision has led to the development of powerful pre-trained models that can be effectively leveraged for a variety of tasks with minimal fine-tuning (Zhai et al. 2022).

In this work, I utilize two such pre-trained models: GroundingDINO (Liu et al. 2024) for object detection and a data efficient version of the Segment Anything Model (SAM) (Kirillov et al. 2022; Chen et al. 2024) for instance segmentation. To provide an accessible entry point for ecologists and evolutionary biologists, I have implemented this deep learning-powered pipeline as both a set of Python scripts and an R package called SegColR. The R framework gives users who are already familiar with the R programming language an easier integration with their existing workflows and analytical tools. Additionally, the Python scripts are available for those researchers who may prefer a more customizable, low-level approach to image processing and analysis. The base SegColR package performs object detection and instance segmentation allowing for individual instances of focal taxa to be extracted from an image. These individual instances then allow for the extraction of color pixel-by-pixel. I demonstrate the SegColR workflow for several examples, showing also how one would assess the quality of the segmentation and color extraction. Finally, I outline potential pitfalls researchers may experience when using this and other automatic object detection and segmentation software.

## 2 Methods and Results

### 2.1 Object detection and instance segmentation

Object detection and instance segmentation are fundamental tasks in computer vision (Chollet 2021). The goal of object detection is to draw bounding boxes around particular objects of interest and associate the bounding box with a particular class for a given image. However, most object detection algorithms are limited to a pre-determined set of classes. This is problematic for biological datasets as existing pre-trained models are unlikely to have been trained on all taxonomic groups of interest and adding new classes would require collecting and labeling new data in order to retrain the model. Attempts to address this challenge have focused on combining visual and textual modalities, with Grounding DINO successfully generalizing object detection (Liu et al. 2024). Grounding DINO is a transformed-based architecture that fuses language and vision modalities by linking the closed-set detector, DINO (Zhang et al. 2022), with grounded language-based pre-training (e.g., GLIP; Li et al. 2022). The effect of this design is that Grounding DINO is able to detect arbitrary objects based on diverse text prompts. This model has been extended in several ways, including combining it with the instance segmentation model, SegmentAnything (SAM; Kirillov et al. 2022). SAM, is an image segmentation model trained on the largest segmentation dataset with over 1 billion masks and 11 million images. This allows it to achieve consistently high performance on zero-shot segmentation tasks even when compared to fully supervised models (Kirillov et al. 2022). The combination of Grounding DINO and SAM is called GroundedSAM (Ren. et al 2024) and it uses the bounding box output of Grounding DINO as the input of SAM for high quality instance segmentation. This approach can be further refined by using recently developed efficient versions of SAM such as SlimSAM, which achieve high accuracy while using far less training data (Chen et al. 2024). SlimSAM result in a model a fraction of the size of the original SAM (1.4% of the original parameters) and is ideally suited for biological research as the workflow can be run on moderately powerful personal computers.

### 2.2 Color extraction

The process of object detection and instance segmentation results in a set of masks for each instance of a particular class. Masks are logical matrices that represent the pixels within an image that correspond to a detected object. Each mask also has a confidence score indicating the model’s certainty about the presence and classification of the detected object. When multiple detections are present for a single class, SegColR combines the masks that meet or exceed a user-specified score threshold. This is a quality control step that users are free to adjust based on the particular needs of their project. On one hand, higher confidence thresholds will be more conservative, but result in the inclusion of only the most reliable object detections. On the other hand, low confidence thresholds will include more detections, but there is a greater chance of inaccurate object detections.

Once the final mask for an object is obtained, SegColR extracts basic color information from the pixels in the original image. This is accomplished by converting the pixel values from the RGB color space to the Lab color space (CIE, 1976) and by default applying a k-means clustering algorithm (MacQueen 1967) to identify the dominant colors within the region of interest. If desired, it is possible to avoid k-means clustering by specifying a set of dominant colors which are then used to cluster each pixel based on the minimum euclidean distance between the pixel color and dominant colors within the Lab color space. The resulting dominant colors are characterized by their Lab coordinates and hexadecimal codes. Additional summary statistics, such as the mean and median color, are also calculated by default. More detailed color analysis is left to the user with other R packages providing more in-depth tool-kits once segmentation has been completed (e.g., Van Belleghem et al., 2018; Maia et al., 2019; Weller et al. 2024).

### 2.3 SegColR features and examples

#### 2.3.1 Description of the SegColR pipeline

Using SegColR requires a user to input an image (specified by the path to the image) and a set of labels. The primary function of SegColR, grounded_segmentation_cli (Table 1), then preforms grounded segmentation based on the input image and labels. This function creates a custom command-line interface (CLI) tool which interacts with a Python back-end. The Python back-end then uses the transformers library, which is itself an API to download and train pre-trained models, to call the segmenter and detector models (Wolf et al. 2020). Note that once the desired models are downloaded and necessary libraries installed, the SegColR software can be used entirely offline. Results are saved as a JSON file which can then be read into R using the function load_segmentation_results. Once the grounded segmentation is complete and the results are loaded into R, the segmentation can be plotted using plot_seg_results and a preliminary color analysis can be conducted using process_masks_and_extract_colors (Table 1). Finally, plot_color_info can be used to display several color summaries of the segmented image including dominant colors, mean and median colors, and RGB histograms. This pipeline is designed in the hope that it will be easy to execute and evaluate the resulting segmentation.

**Table 1:**
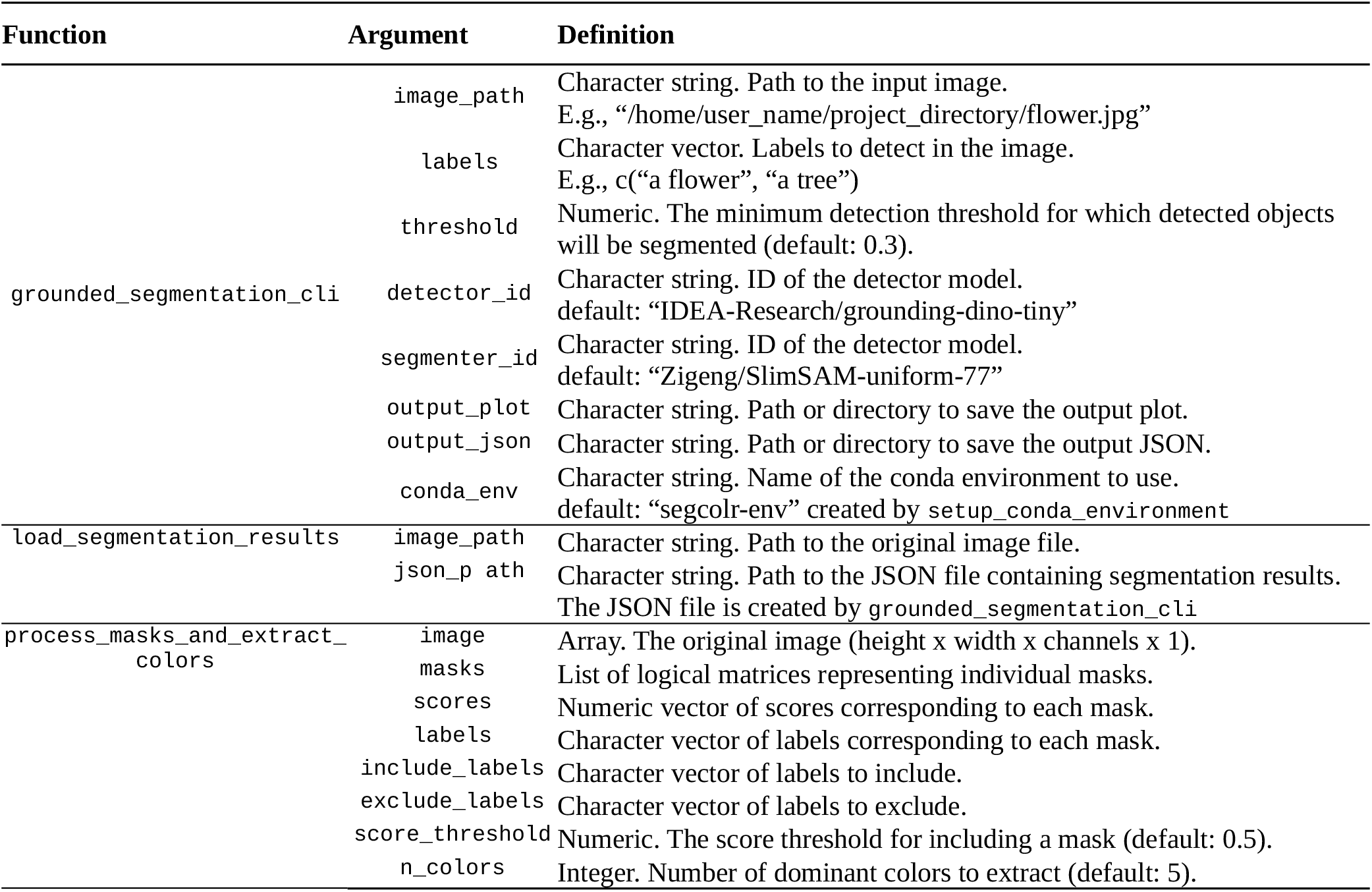
Primary analysis functions.

It is worth noting that a pre-requisite to using SegColR is having all the dependent Python libraries installed for a specified conda environment (conda_env argument of grounded_segmentation_cli). To assist users, I have created the function setup_conda_environment, which will install all the necessary libraries with either exact version numbers known to work with SegColR (env_type= “specific”) or with the minimum constraints on the required libraries (env_type= “general”). I recommend using env_type= “general” in most cases, noting that it takes longer to setup the environment than env_type= “specific” because library versions must be solved to ensure compatibility.

#### 2.3.2 Example 1 – Grounded segmentation and color analysis

This example demonstrates the utility of grounded segmentation in isolating organisms of interest from background information. The subject of this analysis is an Andaman Hind fish, sourced from an iNaturalist observation. Traditional color extraction methods applied to the entire image would yield unusable data due to the background being present in the image. However, by employing grounded segmentation, we can automatically focus on the organism of interest as informed by the user imputed label, significantly improving the accuracy of color extraction. The grounded_segmentation_cli function is utilized to segment the image, with parameters specifying the image path, label, and output paths for JSON data and preliminary plot generation (Table 1).

~~~
ground_results <- grounded_segmentation_cli(
 image_path = “/home/user/images/AndamanHind.jpeg”,
 labels = “a fish.”,
 output_json = “/home/user/output_dir/json/”,
 output_plot = “/home/user/SegColR/extdata/plot/”)
~~~

The output JSON contains information on the score and label for all detected objects as well as the mask for that particular object. The segmentation results are then loaded and plotted using load_segmentation_results and plot_seg_results respectively (Figure 1). Following segmentation, the process_masks_and_extract_colors function is employed to extract color information from the focal organism. This function requires five key arguments, all of which are stored in the output of grounded_segmentation_cli. The results of this color analysis are then visualized using the plot_color_info function, which offers the option to recolor the mask based on the dominant colors extracted (Figure 2). Within R and for ease of use in downstream analysis, both the segmentation results and the color information results are list objects which containing information such as vectors of labels, confidence scores, masks, dominant colors, pixel by pixel coloration, mean and median colors.

~~~
color_results_k <- process_masks_and_extract_colors(
 image = ground_results$image,
 masks = ground_results$mask,
 scores = ground_results$score,
 labels = ground_results$label,
 include_labels = ground_results$label)
~~~

#### 2.3.3 Example 2 – Multiple instances and excluding labels

This example illustrates the capability of the segmentation algorithm to handle images containing multiple instances of a particular label and to exclude overlapping objects and organisms that may interfere with the color analysis. The focus here is on segmenting flowers to extract petal color, in an image that contains multiple flowers and a bee (Figure 3a). The segmentation process assigns scores to each detected instance, allowing for the exclusion of instances below a specified threshold. Setting this threshold to 0.5 results in the retention of two flower instances and one bee instance in the segmented result (Figure 3b). Other than using thresholds, a key feature of the process_masks_and_extract_colors function is its ability to exclude particular labels directly. In cases where multiple organisms are present in a scene, but color is desired from only one of them, specifying each object individually and then focusing the color extraction should produce higher quality results. This functionality is particularly valuable when combined with grounding DINO, as it provides remarkable flexibility in color extraction from non-standard images. In this case, it allows for the removal of the bee instance from the color extraction process (Figure 3c), ensuring that only the colors of the flowers are analyzed.

**Figure 1:**
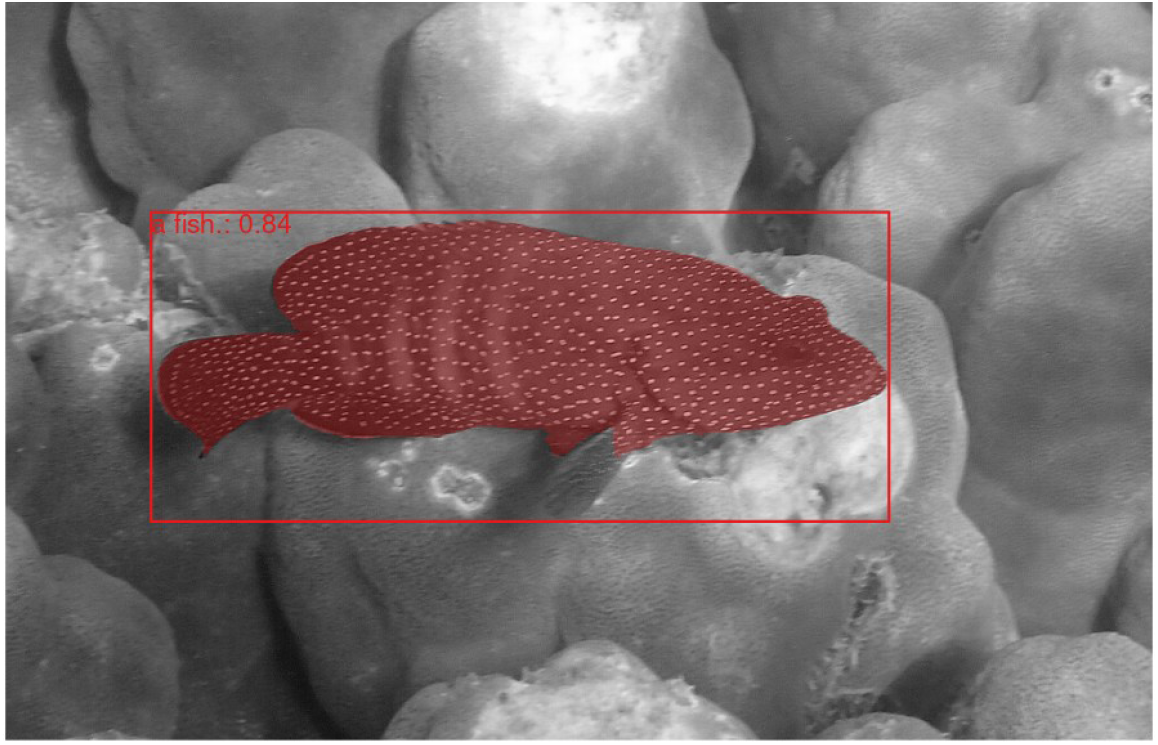
Result of grounded segmentation. Plot produced by plot_seg_results. Confidence score for the object detection (0.84) along with the label (“fish”) are shown in the top left of the bounding box.

**Figure 2:**
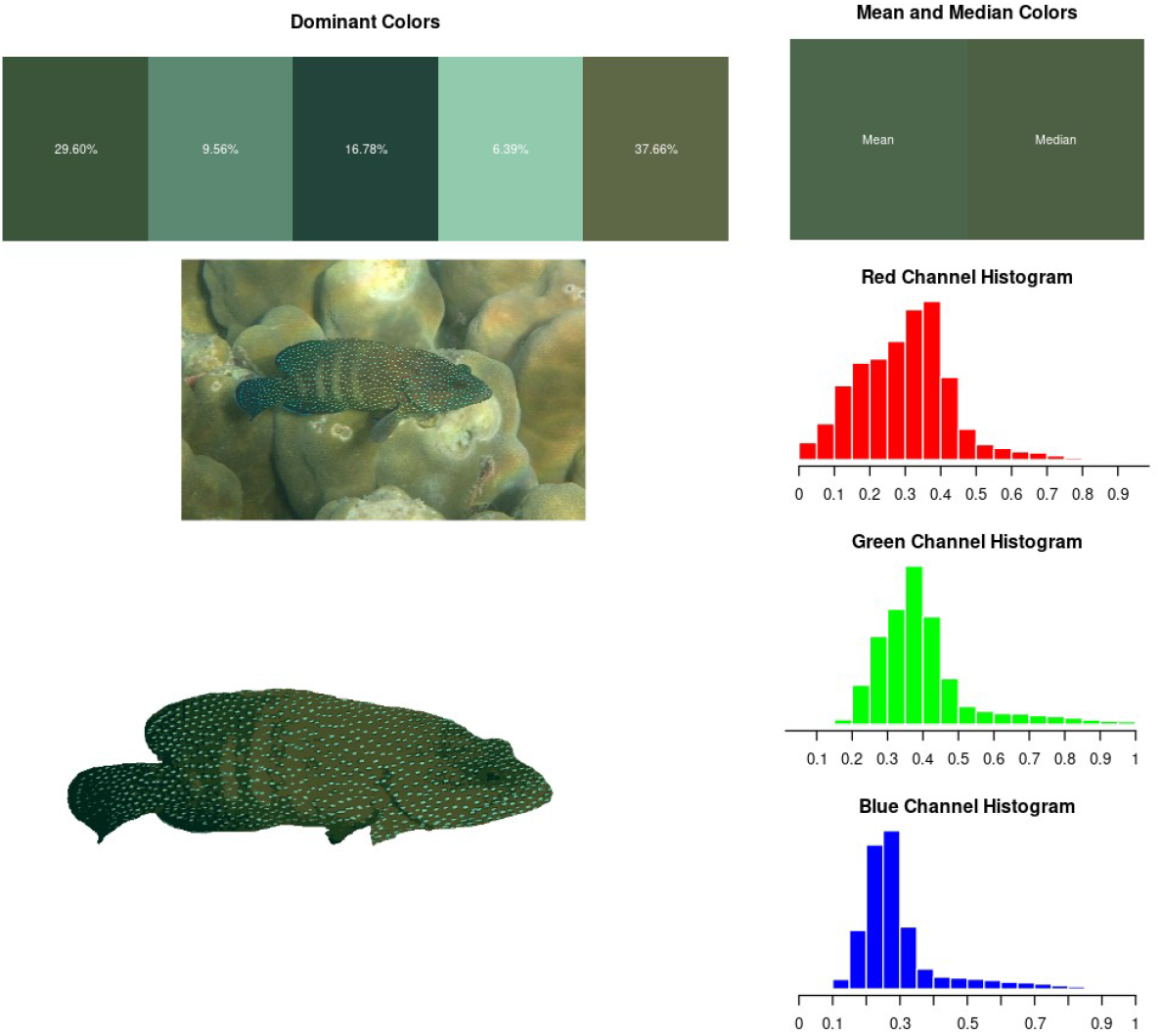
Results of color analysis plotted by plot_color_info. The dominant colors are displayed across the top along with the mean and median colors of the final mask. The right shows the red, green, and blue histogram values between 0 and 1. In the bottom left, the original image and final mask (recolorized based on dominant colors) are shown.

**Figure 3:**
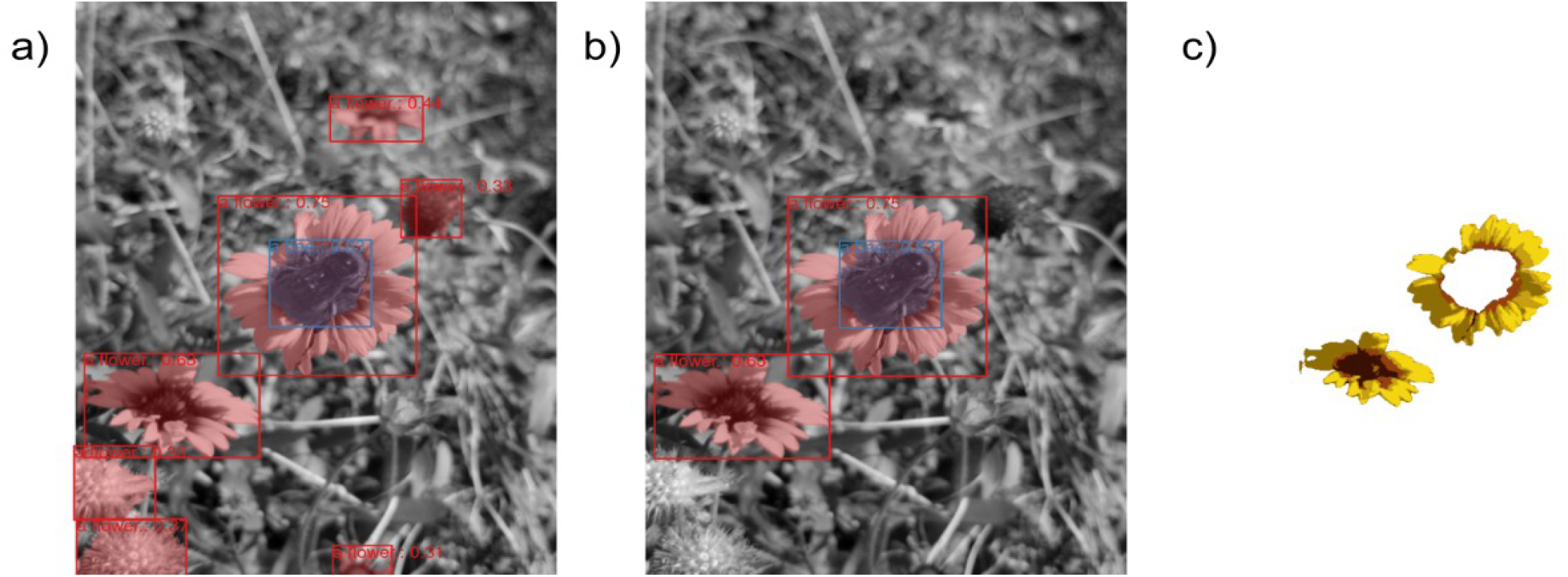
An example illustrating multiple instances of the same object (flower). a) All objects detected. b) Objects above a 0.5 score threshold. c) The final mask for all flowers above a 0.5 confidence threshold and the bee object removed.

~~~
color_results <- process_masks_and_extract_colors(
 image = seg_results$image,
 masks = seg_results$mask,
 scores = seg_results$score,
 labels = c(“a flower.”, “a bee.”),
 include_labels = c(“a flower.”),
 exclude_labels = c(“a bee.”),
 score_threshold = 0.5,
 n_colors = 5
)
~~~

#### 2.3.3 Example 3 – Specifying particular parts of an organism

The final example demonstrates the power of grounding DINO in specifying and isolating particular parts of an organism for focused color analysis. This capability is important when color patterning is localized to specific areas of an organism. The subject of this analysis is a horned bream, where the main color patterning of interest is located on the body. The challenge lies in excluding the fins, which do not contain the coloration of interest. By leveraging the broad textual understanding provided by grounding DINO, we can use text prompts to detect and exclude various fin types from the analysis (Figure 4). This example highlights both the strengths and limitations of the current implementation. While the algorithm successfully identified and excluded the more prominent caudal and pelvic fins, it struggled with the detection of dorsal, anal, and pectoral fins. This suggests that, despite groundingDINO’s generality, fine-tuning the model on domain-specific datasets will still be necessary in some cases.

**Figure 4:**
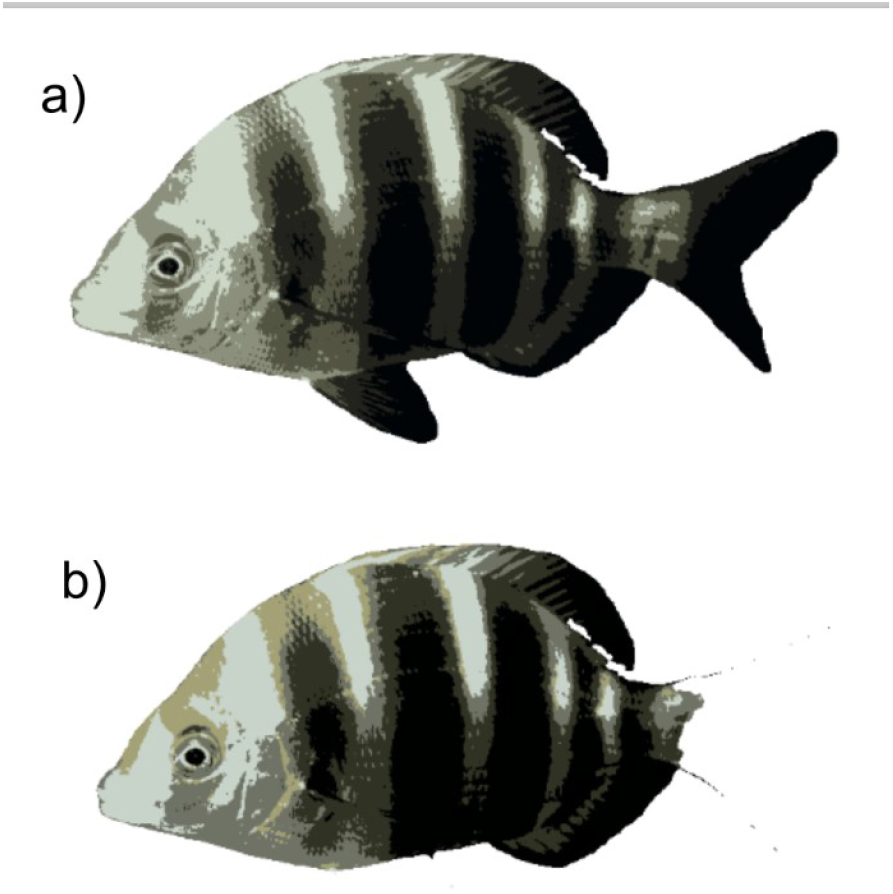
a) The final mask when the entire fish is specified. b) The final mask when fin masks are removed from the fish mask.

~~~
ground_results <- grounded_segmentation_cli(
 image_path = “/home/user/project_file/img.jpg”,
 labebls = c(“a fish.”, “the fins of a fish.”),
 output_json = “/home/user/project_file/json/”,
 output_plot = “/home/user/project_file/plot/”)
~~~

### 2.4 Limitations

The zero-shot object detection capabilities of groundingDINO (Feng et al., 2023) offer significant advantages when processing non-standardized citizen science data, particularly in the context of biodiversity studies. The heterogeneous nature of images on platforms like iNaturalist, where organisms may be positioned anywhere within the frame, camouflaged against diverse backgrounds, or present alongside other species, highlights the value of general text-prompt-based object detection. This approach greatly enhances the accessibility and usability of extensive biodiversity datasets compiled through citizen science initiatives. However, it is crucial to acknowledge that improved object detection algorithms cannot address all inherent biases present in citizen science data. A primary concern is the inconsistency in lighting conditions across images, which can lead to shadow-induced distortions in color values (Szeliski, 2022). While some shadow removal techniques based on luminescence values have been developed (e.g., Murali & Govindan, 2013), their efficacy when applied to the diverse lighting conditions encountered in iNaturalist images has been limited. One potential approach to mitigate these lighting-related issues involves the use of pre-specified color palettes. Within the SegColR framework, this can be implemented through the custom_colors argument in the process_masks_and_extract_colors function. This method clusters pixel colors based on their distance to predefined custom colors rather than employing k-means clustering, potentially offering more consistent results across varying lighting conditions.

Beyond these technical limitations, citizen science datasets are subject to broader constraints that merit consideration. A significant issue is the presence of sampling effort biases (Dickinson et al., 2010). The majority of data is generated from North America, particularly the United States, despite the fact that global biodiversity is concentrated in tropical regions, which remain critically undersampled (e.g. Vasconcelos, 2023). While initiatives to increase sampling efforts in tropical areas exist, current datasets exhibit substantial geographical biases that must be accounted for in analyses utilizing citizen science data (Ward, 2014). Another limitation specific to color analysis is the discrepancy between human color perception (and standard camera capabilities) and the color perception of various organisms (Endler & Mielke, 2005). Many species perceive colors in ways that differ significantly from human vision, often requiring specialized equipment to capture color accurately. The widespread adoption of such specialized imaging technology among citizen scientists contributing to platforms like iNaturalist is unlikely, potentially limiting analyses to the human-perceptible color space. However, it is worth noting that images captured with specialized equipment could still be processed and analyzed using tools like SegColR, should they become available.

## 3 Discussion

The proliferation of citizen science initiatives has led to an unprecedented accumulation of biodiversity data, offering researchers a vast repository of information. However, the heterogeneous nature of these datasets presents significant challenges for standardization and analysis. Recent advancements in computer vision technologies have emerged as a promising solution to address these complexities, offering remarkable flexibility in data processing and quantification. The application of deep-learning to tasks within ecology and evolutionary biology is not new, however, to-date, most applications have focused on models trained from scratch (e.g., Weaver and Smith 2023). A more powerful, accessible, and environmentally responsible way forward for academics may be the utilization of large pre-trained models. These models are trained on extensive datasets which few academic collaborations are able to match (e.g., the billion masks used to train SAM; Krillov et al., 2023). Fortunately, many of the underlying representations within the hidden layers of the deep-learning models are still useful in biological studies and through the use of fine-tuning, can be readily adapted to even the most obscure taxonomic groups.

The implementation of deep learning tools in R is an important step towards making these techniques widely available to biologists. R is one of the most widely used programming language for academic ecologists and evolutionary biologists, but most deep-learning developments take place in Python. Furthermore, the use of light-weight models, such as slimSAM (Chen 2024), allows for advanced deep-learning models to be run even on moderately powerful personal computers. By enhancing the accessibility of these tools, biologists can gain access to an increasing number of data sources. Nonetheless the automation provided by computer vision techniques will need to be balanced with careful verification of the results. Performance is not guaranteed to be the same across all images (see example 2.3.3) and increasing the accuracy of these models on diverse taxonomic groups will likely require some amount of additional data collection and fine-tuning. Furthermore, while SegColR allows for a more automated collection of citizen science data, it cannot address all inherent limitations of this data source. Researchers must remain cognizant of lighting inconsistencies and sampling biases when interpreting results derived from these datasets.

## 4 Conclusion

Color is a crucial phenotype for many organisms, however gathering large interspecific datasets has proven difficult without some amount of automation. Here I have introduced SegColR, an R package which utilizes an underlying Python framework to automatically detect and segment organisms within citizen science photos. Using the pre-trained models groundingDINO and slimSAM, SegColR is able to preform object detection and segmentation without training for specific taxa. Using a computer vision pipeline for automated color extraction is just one example of how deep learning techniques can be used to quantify large-scale bio-diversity datasets. SegColR and other deep-learning software offer many exciting possibilities, but still necessitate careful consideration of the data produced. Researchers must reconcile the fact that though these tools are able to automatically extract vast quantities of data, high quality datasets will only be created if they are rigorously evaluated. As deep-learning techniques continue to integrate with ecology and evolutionary biology, it is important that they are used to complement traditional methods, rather than replace them.

